# The Genomic Prediction of Disease: Example of type 2 diabetes (T2D)

**DOI:** 10.1101/285783

**Authors:** Lawrence Sirovich

## Abstract

Application of concepts from information theory have revealed new features of Single Nucleotide Polymorphism (SNP) organization.. These features lead to effective classifiers by which to distinguish genomic sequences of contrasting phenotypes; as in case/control cohorts.

When applied to a disease/control database, a disease classifier results; a parallel analysis leads to the determination of a wellness classifier. The classifiers have non-intersecting loci, and each involves roughly 100 alleles.

The effectiveness of this framework is illustrated by application to adult onset, type 2, diabetes (T2D), as represented in the Wellcome Trust ((WT) Case/Control database.

Simultaneous use of the two classifiers on the WT database leads to successful prediction of disease versus wellness; to the extent that near certain genomic forecasting is achieved.

This framework gives a resolution to the oft posed uncertainty: “Where is the missing heritability?”

Application of both classifiers on two additional T2D databases produced informative consequences.

A fully independent, compelling, confirmation of the present results is obtained by means of the machine learning algorithm, Random Forests.

The analytical model presented here is generalizable to other diseases.

**One Sentence Summary:** Discovery of intrinsic chromosomal SNP organizations leads to near certain genomic disease prediction.

## 1. Introduction

The connection of genomics to human diseases has long been under investigation. In pursuing the contrast between disease and wellness, one compares statistical measures of both disease and control populations, under the reasonable hypotheses that statistical dissimilarities will reveal risk loci. The conventional approach in identifying potential dissimilarities follows from odds-ratios considerations for disease versus control populations (1). The availability of quality Genome Wide Association Studies (GWAS) databases has produced an abundance of loci deemed to be high-value, but this circumstance is accompanied by qualms about low population penetration, i.e., low incidence of the high-value loci in the data (2); and hence a weak capacity for disease identification. Family histories, which play a role in GWAS case/control selection, appear to be more informative than genomics in disease identification (3). This state of affairs has led to a collection of publications raising the question “Where is the missing heritability?”(4-6). Moreover, the abundance of disease risk loci coupled with the lack of power in accounting for disease has aroused skepticism in regard to the GWAS effort itself (7-12).

An effort by the author to apply novel mathematical tools to the genomic prediction problem produced disappointing results (13). Further investigation revealed this to be a consequence of an inherent bias in the odds-ratio; low penetration loci are unduly emphasized. Improvement came with the use of ideas from information theory (14), which led to an unbiased selection criterion referred to as incremental information (15). Additionally, this process revealed the existence of sets of genomic loci which have the property that disease and control sequences project into well-separated clusters (15).

Analysis of the WT genomic sequences indicates that, along each chromosome, high-value loci tend to arise in loose clusters, and that this clustering is far more pronounced in the disease set then in the control set.

This feature recalls the frequent observation that complex diseases arise from large collections of weakly contributing loci (7, 9, 11, 16), which leads to the construction of classifiers, which permit the inclusion of many genomic loci.

The methodologies which lead to these results differ from common practice (1). Nevertheless, they follow from simple arguments and plausible premises. A summary of methods in Section 7 is presented in this spirit.

## 2. Risk Allele Linkage

A well-studied disease/control model is Type 2 diabetes, T2D, (17-22). This will serve as the illustrative example case of the present analysis, and the WT T2D database (22) will be the principal focus.

The WT database can be regarded as comprised of two matrices, *D* and *C*, that describe the disease and control cohorts. *D* has 1999, and *C*, 3005 rows; each row is the genomic sample of an individual; both matrices have 990,950 common allele columns. Instead of treating the full database it will be simpler and more revealing to examine a subset of high information alleles chosen so that the incremental information, *IF* (7.3), at each such allele, satisfies the criterion

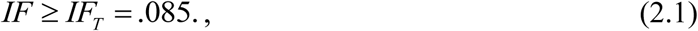

This results in the selection of 1007 high-value alleles. The reason for the criterion, (2.1), will appear later.

The disease state can be associated with the rare variant allele, the *rare variant hypothesis*, see Methods, section 7. The data show that the rare variant alleles have a 50% higher presence in the disease population than in the control set. This observation enters in the derivation of incremental information as a selection procedure (15).

### Linkage

The restriction of matrices *D* and *C* to the 1007 selected alleles that satisfy (2.1) is denoted by *D°* and *C°*.

It is hypothesized that consecutive rare variants in sequences (rows of *D°*) is an indication of the disease *mechanism*; hence the control population should show less of this *linkag*e. To test this hypothesis a *linkage* matrix *L*^*D*^ for *D°* is constructed, one chromosome at a time, as follows: If an entry of *D*^*°*^ is the appropriate rare variant entry then *L*^*D*^ at this location is given the value of 3 if it is both preceded and succeeded by the appropriate rare variants; if only one of these is present then *L*^*D*^ is given an entry of 2; if there are no neighboring rare variants matrix entry is 1; all other entries of *L*^*D*^ equal 0. This nearest neighbor approximation is adequate for our purposes. Although this may seem related to the haploid assumption, it is not, since successively loci are typically megabases apart.

The mean linkage value of *L*^*D*^ at the *j*^*th*^ allele, or simply the *j*^*th*^ linkage, is defined by its mean value over the *j*^*th*^ column,

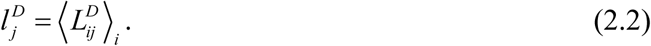

Similarly, 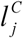 is obtained from the linkage matrix of *C°, L*^*C*^. The *j*^*th*^ linkage ratio at an allele is defined by

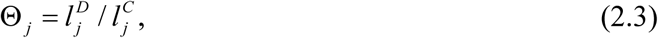

and based on the data we find the mean

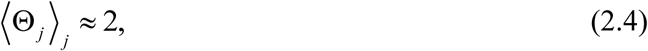

which confirms the hypothesis, that disease sequences exhibit significantly higher linkage than do control sequences; this furnishes one more tool for distinguishing disease versus control.

## 3. Classifiers

The purpose of this section is to introduce the concept of a classifier, as a determinant of phenotype. Within the subset of the alleles that satisfy (2.1), a subset of alleles, *Al*, a vector of locations, is chosen. The corresponding vector of risk symbols at the loci *Al*, is denoted by*W*, and the classifier defined by,

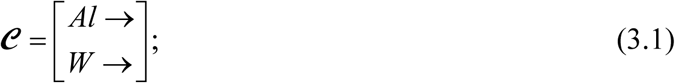

a two rowed matrix of locations and symbols. *W* is a *word* in the general sense. Any genomic sequence can be viewed as a *word* defined on the classifier loci *Al*, which word can then be compared (by the Hamming distance) with the classifier word *W*, itself.

For this purpose consider a purported sequence, *s*, then its classifier score is

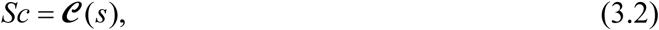

alculated by the calculated agreement with *W*.

If the phenotype is disease, the disease classifier is denoted by

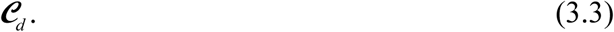

Therefore for some arbitrary sequence, s, the score of this classifier is expressed as

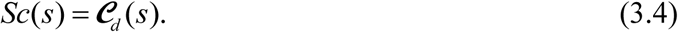

An underlying symmetry of the formulation suggests that one can also contrast the control cohort with respect to the disease cohort. The goal is then a *protective* or *wellness* classifier which will be denoted by

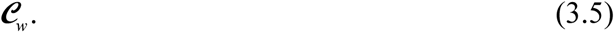

In simple terms, (3.5) is obtained by mimicking the disease classifier analysis under the interchange of roles for the disease and control populations. The two resulting allele sets are non-intersecting, i.e. intersection of the alleles of ***𝒞***_*d*_ and ***𝒞***_*w*_ is empty, since they are chosen by a complementary incremental information, *IF*, criterion (15).

Conceptually, the protective set is composed of loci of beneficial mutations, disease antagonists; loci that represent potential resistance to the disease. This is speculative and the terminology is regarded as metaphorical; however, such mutations have been reported in the literature (23).

The two classifiers, (3.3) and (3.5), can be used in a general classification procedure as follows: For an arbitrary sequence s, a net differential score is calculated by,

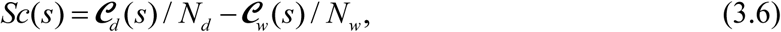

to serve as a phenotypical determinant of the sequence *s. N*_*d*_ and *N*_*w*_ are the corresponding classifier word lengths by which to normalize (3.6).

If *Sc(s)* is positive *s* is deemed to be a disease sequence and if negative deemed to be a control (protective) sequence. *The state of any putative sequence is determined by which classifier, disease or wellness, better fits the sequence.*

It is at least of passing interest to point out that the disease and control classifier scores can be shown to be well fit by Gaussian probability distributions (15), the overlap of these Gaussians can be measured by

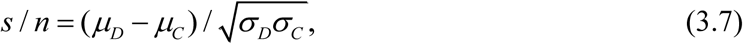

where *µ* and *σ* refer to the population mean and standard deviation values, respectively. This measure may be rightly regarded as a signal-to-noise ratio, as denoted.

A simple distance between clusters is given by (15),

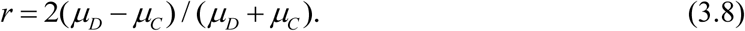

## 4. Verification and Prediction

The pair of parameters, *IF* and Θ, are key to the discussion. Incremental information, *IF*, can be regarded as the alternative to the commonly used *odds ratio.* The selection of threshold criterion *IF* > .085, (2.1), prunes the number of possible alleles by a drop in their count from *O*(10^6^) to *O*(10^3^).

A standard training/test paradigm will be applied to assess the parameter space with the goal of choosing a sensible parameter pair. Briefly, we survey the parameter rectangle

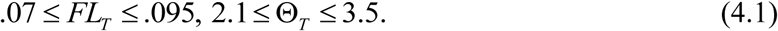

In particular classifier size, signal-to-noise, cluster separation and success rates are monitored over this range. This strategy was carried out over many random trials of the train/test protocol with a 98%/2% splitting, which should give a sense of how well classification is achieved on novel sequences. The principal objective of this search will be the determination of minimal sized classifiers. The search led to the choice of a threshold pair in the neighborhood of the lower right-hand corner of the parameter rectangle (4.1).

One piece of this pair is the incremental information choice, (2.1); in the same spirit the remaining parameter, the linkage threshold is chosen to be,

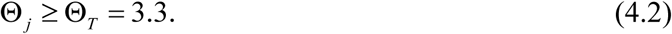

To get a sense of the reach of the classifiers a variety of partitions were considered.

In all cases for which the training set was greater than 90% of the database, successful prediction exceeded 98 %; when the fraction was 50%, the success rate exceeded 88%; and even at 25%, the success rate was almost 70%.

A 90% training 10% test division was performed 50 times, so that the possibility of a sequence not being tested was less than 1%. The result was a success rate of

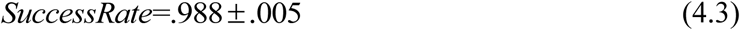

In this instance the derived disease classifier, ***𝒞*** _*d*_, is composed of 97 alleles and the wellness classifier, ***𝒞*** _*w*_, of 99 alleles.

Since (4.3) might well be regarded as *too good to be true*, further explorations were undertaken. A return to the 98%/2% splitting, showing how well classifiers do with novel sequences was performed 250 times, again to reduce the chance that a sequence would not be tested to less than 1%, and the result was

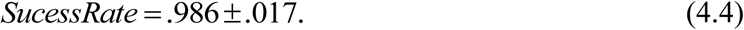

Finally, 250 randomly chosen 99.5%/.5% divisions were performed. Each test set then consisted of roughly 10 disease and 15 control sequences. This yielded

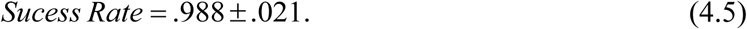

In this instance 186 out of the 250 trials gave perfect prediction, and 57 had one error and the remaining 7 trials two errors.

Although, the success rate appears to be stationary, at roughly 99%, the signal-to-noise ratio, *s/n* and the cluster separation, *r*, show small improvement in passing from (4.3) to (4.5). More important is the decline in disease classifier size from 97 to 89 alleles.

Another, completely independent, confirmation of these results will be presented in the next section.

For publications which present predictive models, it has become routine to perform some sort of data randomization in order to deal with the possibility of overfitting. Since the above procedures produced successful classifiers that have fewer than 200 alleles, out of a total of million alleles, and for over 5000 sequences, there is little likelihood of overfitting. Nevertheless, a full randomization of disease and control data by shuffling all sequences was undertaken, and led to no predictive success except for rare and isolated instances due to pure chance.

## 5. Comparisons

### Other Databases

Two additional T2D databases were available for testing. The testing was performed in two ways: (1) The above classifiers, ***𝒞*** _*d*_, *and* ***𝒞*** _*w*_, were transformed into an invariant form that could be applied to any T2D GWAS database and on this basis success rates could be tested; (2) A de novo application of the above framework is applied to each GWAS database, and tested by the training/test procedure used above.

#### Fusion T2D (21)

A prior investigation (13, 15), was based on the Fusion T2D database. This database is composed of 272,423 SNPs, and contains 919 disease and 787 control sequences.

The translated form of ***𝒞*** _*d*_ was empty, and that of ***𝒞*** _*w*_ only contained seven alleles, thus rendering the invariant form useless for the Fusion database. In this regard it should be noted that the intersection of the full SNP sets for the Fusion and Wellcome Trust databases numbered 45,311, less than 10% of the Wellcome Trust database.

A de novo application of the present framework produced the *Success fraction =.5001±.0389;* regarded as pure chance. One might conclude from this that the gene array selected for the Wellcome Trust database was better suited for revealing the genomic basis of T2D.

#### Starr County T2D (18)

In this case the invariant forms of, ***𝒞*** _*d*_, *and* ***𝒞*** _*w*_, contained 82 and 88 alleles respectively, however, the success rate was a very modest 51%. A de novo treatment of this database produced a *Success fraction =.5340 ±. .0264.*

The reason for the robust size of the transformed classifiers can be traced to the fact that the Starr County study used the Affymetrix 6 gene array which contained the Affymetrix 500 K Snps used in the WT study. However, the key observation is that the Starr County disease and control populations are comprised of Mexican Americans, and therefore possess a significant element of the Native American genome. This implies a very wide gap in ancestries for the WT and Starr databases; possible conclusions are that the WT Snps were poorly suited for the Starr County subjects, and that the Starr County gene array did not go deep enough in the set of SNPs to account for their ancestry, see Discussion.

#### ¶Random Forests

Random Forests (RF), a machine learning algorithm has recently been applied to the WT T2D database (24, 25).

The following brief description should give reader a sense of this method. This method is built on random data sampling, with replacement; building learning trees which pivot at suitable features and which in conclusion determine prediction on the basis of a vote by the “trees of the forest”. Instead of classifiers the result of RF is a tree structure that can be applied to test sequences to determine success rate on the basis of a ballot. Both cited references have notable results. In both instances some standard data filtering (15) is used to determine the features to seed the RF algorithm; (24) also contains a tweaking of the standard RF algorithm. Both investigations indicate success prediction level that approaches 90%.

In order to compare the present deliberations with RF, the alleles of both disease and wellness classifiers were combined and introduced these as features to seed the RF algorithm. The resulting success rate was close to the present 99% range. Thus RF prediction rose to a significantly higher level. This calculation serves as a fully independent verification of the present deliberations and reveals the RF algorithm as a tool for GWAS investigation.

## 6. Discussion

Examination of the disease and wellness classifier loci, (3.3) and (3.5), reveals that they are mostly associated with introns and therefore their relation to the disease mechanism is ambiguous. However, since they foreshadow the disease in such a compelling manner they likely play a, still to be determined, role.

In contrast to the highly specific loci of Mendelian diseases, T2D risk loci were found on almost all chromosomes, 22 and 23 being the exceptions. Another departure from the Mendelian case is that disease and control sequences share the same allele risk loci, but with higher frequency in the disease population. This commonality of allele symbols occurring in the disease and control population has been observed in the literature (26-29) and has been termed the common *disease/deviant* hypothesis.

Neel (30) suggested that T2D, as a disease, might be due to a past mutation, which for nutritional reasons was once advantageous, but has since been “rendered detrimental by progress” but not so detrimental as to become eliminated through natural selection. One can imagine that over the course of the tens of millions of years of primate evolution, the introduction of such mutations are a recurring variation on a theme; disease loci appear on almost all chromosomes. One can also imagine that in the course of evolution, mutations antagonistic to a disease might also arise. (A recent paper reports on a beneficial mutation for Alzheimer’s disease (23).) The ‘wellness’ classifier was introduced in this spirit, but served mainly as a useful technical device for identifying the disease versus control state of a putative sequence. Possibly, the wellness loci as found here might serve as an entry point in the search for beneficial SNPs for T2D.

In regard to the issue of “missing heritability” as raised in the Introduction, it was pointed out in (5) that based on odds ratios selection the overwhelming number of high risk loci showed “small increments in risks, *rr*,

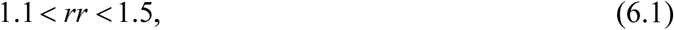

which only explain a small portion of heritability”. Here *increments* refer to the risk ratio or relative risk, *rr*, the ratio of disease/control of nucleotide probabilities. By contrast, for the loci of the disease classifier obtained here, under incremental information selection, we obtain risk ratios satisfying,

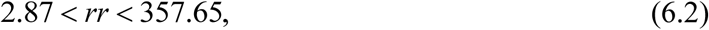

orders of magnitude larger than that obtained under the odds ratio criterion, (6.1), which eliminates the issue of penetration.

In another vein we mention two recent studies, (19, 20) that contain a combined list of 95 SNPs that are regarded as possessing high susceptibility to T2D (on the basis of odds ratio selection). The intersection of the 95 SNPs with both classifiers derived here is empty. We also find that only 29 of the *high susceptibility* SNP list appear in the Wellcome Trust T2D collection. Investigation of the 29 SNPs reveals that these all show small penetration of the sort indicated in (6.1). Additionally, these loci have low incremental information, which explains the *empty* intersection just mentioned.

As indicated in (3) the notion of ‘heritability” has been loosely defined, which accounts for some imprecision in usage. Largely, the term “missing heritability” is used to describe the disappointing identification of disease on the basis of loci deemed to be of high-value. Replacement of the odds ratio by incremental information in the selection of high-value alleles brings with it high penetration, (6.2), and demonstrates that disease prediction can be performed with near certainty, and the elimination of “missing heritability”. This is in no small way due to two unique features of the Wellcome Trust database.

Seven different diseases figured in this study. Compared to a single disease study this entailed a sevenfold increase in the search range for relevant SNPs to serve as *reporters* of the gene array. Secondly, the control cohort was chosen to be free of all seven diseases; which furnishes a near ideal baseline for the contrast of disease versus control. A comparative study of several databases testing their incremental information range was performed and convincingly demonstrated that the Wellcome Trust database indeed provided the widest range of Incremental Information values.

Future GWAS databases investigators might wisely follow the lead of the Wellcome Trust initiative to achieve similar levels of range and precision. It is also suggested that the use of incremental information be adopted in the preliminary studies that usually precede reporter selection for a gene array.

## 7. Methods Summaries

### A Data Description

The principal content of a GWAS database, lies in its disease and control matrices, *D* and *C*. The matrix columns of these are the SNPs which appear in chromosomal order, and then within each chromosome in increasing distance from a fiducial point.. The two alleles of each SNP are taken to be universal, and a priori known. Each SNP occupies two successive columns, the total number of columns is twice the number of SNPs.

Each row of a matrices contains the genomic content of a single individual; clinically deemed to be of the disease or control cohort. Matrix entries are alleles, from [*A,C,G,T*], but for manipulation are aliased by the integer symbols, [1,2,3,4]. *This is not a digitalization*.

### Outline of Methods

I. *Alleles*: The known pair of alleles, of a SNP are specified in a file of the GWAS database. For uniqueness in representation the alleles of a SNP are placed in anti-integer order. E.g. if the SNP pair in a sequence is (2, 4) it appears as (4, 2). Since the allele pair is known, the representation is further reduced by expressing it as (2, 1). This states that both the high integer and low integer alleles of the SNP both appear. Since the correct symbols are known this, this can be done in general and entails no loss of information. For the example SNP there are the three possibilities,

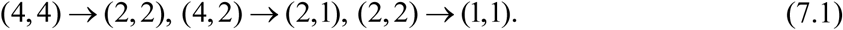

A SNP will always have the designated first and second symbols, 2 and 1, respectively. The further transformation from (2, 1)→ (1, 0) reduces all sequences to binary form.
II. *Probabilities*: For any data matrix the probability, based on frequency, of symbol 2 at the first allele is denoted by *p*_1_ (2), and from this *p*_1_ (1) = 1− *p*_1_ (2). Similarly, for *p*_2_ (1) & and *p*_2_ (2) = 1− *p*_2_ (1).
III. *Information:* According to Shannon (14, 31), the information contained in the first alleles, *I*_*1*_, is given by

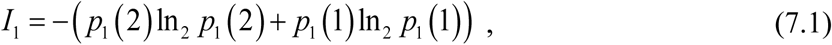

with an analogous form for *I*_*2*_.
IV. *Allele symbols are binomially distributed*: Since each allele has one of two symbols and under the assumption of independent sequences the distribution is rigorously binomial. This is confirmed by the data.
V. *Rare allele hypothesis*: Denote by *Q*_*d*_ and *Q*_*c*_ the minor or rare allele probabilities of the disease and control populations at a specific locus. For a disease such as T2D, since the majority of human beings are without the disease, it might be inferred that *Q*_*d*_ *< .5* at a disease locus, and thus that *risk alleles* should satisfy

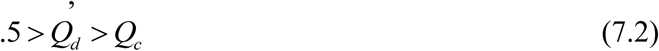

as a guide for selecting candidate loci. While this argument appears reasonable, it might not be compelling. Nevertheless, the data suggests that it is true, cf. (32).
VI. *Incremental Information:* Customarily, odds-ratios are used to locate candidate risk loci, however as shown in (15), this criterion is biased in favor of low probabilities which can be misleading. This criterion is here replaced by a criterion based on information theory, termed incremental information,

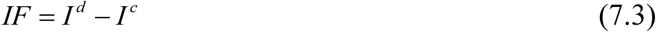

where *I* ^*d*^ and *I* ^*c*^ are disease and control information at the same allele, defined by (7.1). Inspection of (7.1) shows that the role of low probabilities is diminished.

## Acknowledgments

The calculations reported on here were carried out with the aid of Matlab.

Thanks to Cameron Rudd and Avinoam Henig for their help with data preparation. This investigation could not have taken place without the high quality data supplied by the Wellcome Trust, which I gratefully acknowledge. Partial support for the present research came from an award from the Robertson Fund under the auspices Rockefeller University.

Finally warm thanks to Mitchell Feigenbaum for affording me the hospitality of Rockefeller University.

